# Mechanistic variability in corneal nerve recovery linked to injury type

**DOI:** 10.1101/2025.03.18.643516

**Authors:** Léna Meneux, Sarah Pernot, Nadège Feret, Melissa Girard, Alicia Caballero Megido, Naima Nhiri, Elea Miessen, Eric Jacquet, Frederic Michon

## Abstract

The cornea, a transparent tissue covering the eye, is essential for clear vision and represents the most densely innervated tissue in the body. Its extensive sensory innervation provides both sensory perception and crucial trophic support, maintaining corneal health and integrity. Disruption of corneal innervation leads to neurotrophic keratitis (NK), a pathological condition caused by ocular injury, surgical procedures, or underlying diseases. The limited understanding of NK’s pathophysiological mechanisms has hindered the development of innovative therapeutic approaches. In this study, we comparatively analyzed corneal innervation morphogenesis and regeneration across two clinically relevant injury models, highlighting both commonalities and differences among these contexts. Our results demonstrate that corneal nerve morphogenesis and maturation span approximately 13 weeks, from embryonic day 12 (E12) through three months of age. Additionally, we observed that the specification of distinct nerve fiber types coincided temporally with a significant enhancement in corneal sensitivity. Furthermore, we found that the type of innervation loss—either via axotomy or abrasion—differentially affected corneal sensitivity and epithelial cell homeostasis. Importantly, the regeneration mechanisms following injury were also distinctly dependent on the type of nerve damage sustained. Collectively, these findings underscore both the shared characteristics and unique aspects inherent in each NK model, highlighting the necessity for tailored therapeutic strategies specific to individual patterns of corneal innervation disruption.

## Introduction

The unique structural organization of the cornea is essential for maintaining its transparency and mechanical robustness. Representing approximately 75% of the refractive power of the eye, the cornea also acts as a protective barrier against external injuries^1^. Any disturbance to its structural integrity can severely compromise vision, potentially leading to blindness. Structurally, the cornea consists primarily of the stroma, accounting for about 90% of its total thickness, which is predominantly composed of extracellular matrix components and sparsely distributed keratocytes. The anterior stroma is covered by a continually renewing epithelium composed of five to seven layers of cells, while the posterior surface is lined with a monolayer of endothelial cells^2^. Corneal transparency is heavily reliant on meticulous maintenance of tissue homeostasis. This homeostatic balance is sustained through a specialized corneal microenvironment designed to minimize potential damage and maintain structural integrity. Key elements of this microenvironment include the tear film secreted by lacrimal glands, the corneal epithelium, and the dense network of corneal nerves, which collectively provide a rich blend of trophic factors essential for corneal health^3–5^.

The cornea is notably one of the most densely innervated tissues in the body, predominantly innervated by sensory nerve fibers (C and Aδ types) originating from the ophthalmic branch of the trigeminal ganglion^6^. During mouse embryonic development, sensory fibers extend from the trigeminal ganglion along the optic cup, ultimately penetrating the corneal stroma via four distinct quadrants: dorsal-nasal (DN), dorsal-temporal (DT), ventral-nasal (VN), and ventral-temporal (VT)^7^. These nerve fibers extensively branch across the corneal surface to establish the subepithelial plexus. Subsequently, these fibers make a perpendicular turn, piercing Bowman’s membrane and forming the subbasal plexus, recognizable by its characteristic central vortex. Finally, the nerve fibers extend another 90° turn to terminate as free nerve endings at the epithelial surface^6,8,9^.

In the mature cornea, several classes of nerve fibers have been identified, distinguished by their functional and electrophysiological characteristics^10^. Cold receptors respond to temperature changes and express the transient receptor potential cation channel subfamily M member 8 (TRPM8), while mechano-nociceptors respond primarily to mechanical stimuli through activation of the Piezo2 channel. The most abundant fibers, polymodal nociceptors, can respond to mechanical, thermal, and chemical stimuli. Together, these nerve fibers play critical roles in protective mechanisms, triggering reflexes such as blinking, promoting tear secretion, and releasing trophic factors necessary for maintaining corneal integrity^11,12^. Corneal innervation disruption results in neurotrophic keratopathy, a degenerative pathology characterized by nerve loss, reduced corneal sensitivity, and compromised trophic functions^13,14^. Clinically, neurotrophic keratopathy progresses through three distinct stages: initially manifesting as punctate keratopathy and reduced tear stability, advancing to persistent epithelial defects with associated loose epithelium and stromal edema, and culminating in severe ulceration, stromal melting, and potential perforation^15^. Common etiologies include herpetic infections, trauma, chronic conditions such as diabetes, and surgical interventions^13,14^. Treatment for neurotrophic keratopathy typically begins with topical application of therapeutic eye drops formulated to stimulate healing^14,16–19^. In cases where topical therapies prove ineffective, surgical interventions such as amniotic membrane transplantation^20–22^, keratoplasty^14,23,24^, or nerve grafting (neurotization)^25,26^ may be considered. However, treatment failure and graft rejection remain significant challenges despite repeated interventions^27,28^.

Given the limited understanding of molecular pathways involved in axon guidance and regenerative growth within the cornea, therapeutic options remain restricted. Our study addresses this knowledge gap by characterizing the developmental patterns of corneal innervation in mouse embryos, focusing on specific mechanisms guiding axon formation and maturation. Additionally, we examine nerve regeneration processes utilizing two surgical mouse models that parallel clinical scenarios such as foreign body injury or lamellar keratoplasty-associated trepanation.

## Results

### Corneal sensory innervation embryonic morphogenesis

Given the crucial roles of the cornea in transparency and mechanical integrity, understanding its innervation during development is essential. Since nerve fiber bundles begin reaching the cornea around embryonic day 11 (E11) in mice, we focused our analysis between E11 and E14 to characterize the establishment and initial patterning of corneal axons (**Figure 1**). To visualize the morphogenesis of corneal innervation, we utilized light-sheet microscopy on transparized embryos and immunolabelling of βIII-tubulin, a pan-neuronal marker. At E11, nerve fiber bundles initiate their trajectory toward the cornea, extending along the optic cup. By E12, these bundles have reached the corneal periphery, where they initially deflect by approximately 90°, remaining peripheral rather than penetrating directly into the cornea. Notably, innervation on the dorsal side of the cornea is consistently delayed compared to other regions. By E14, nerve fiber coverage has expanded significantly, with extensive branching nearly encircling the entire corneal circumference, reminiscent of chicken corneal innervation morphogenesis. Interestingly, fibers appear to exhibit repulsive interactions, avoiding direct contact with one another. These observations align with previous findings but contribute novel, detailed insights into the process.

**Figure 1.**
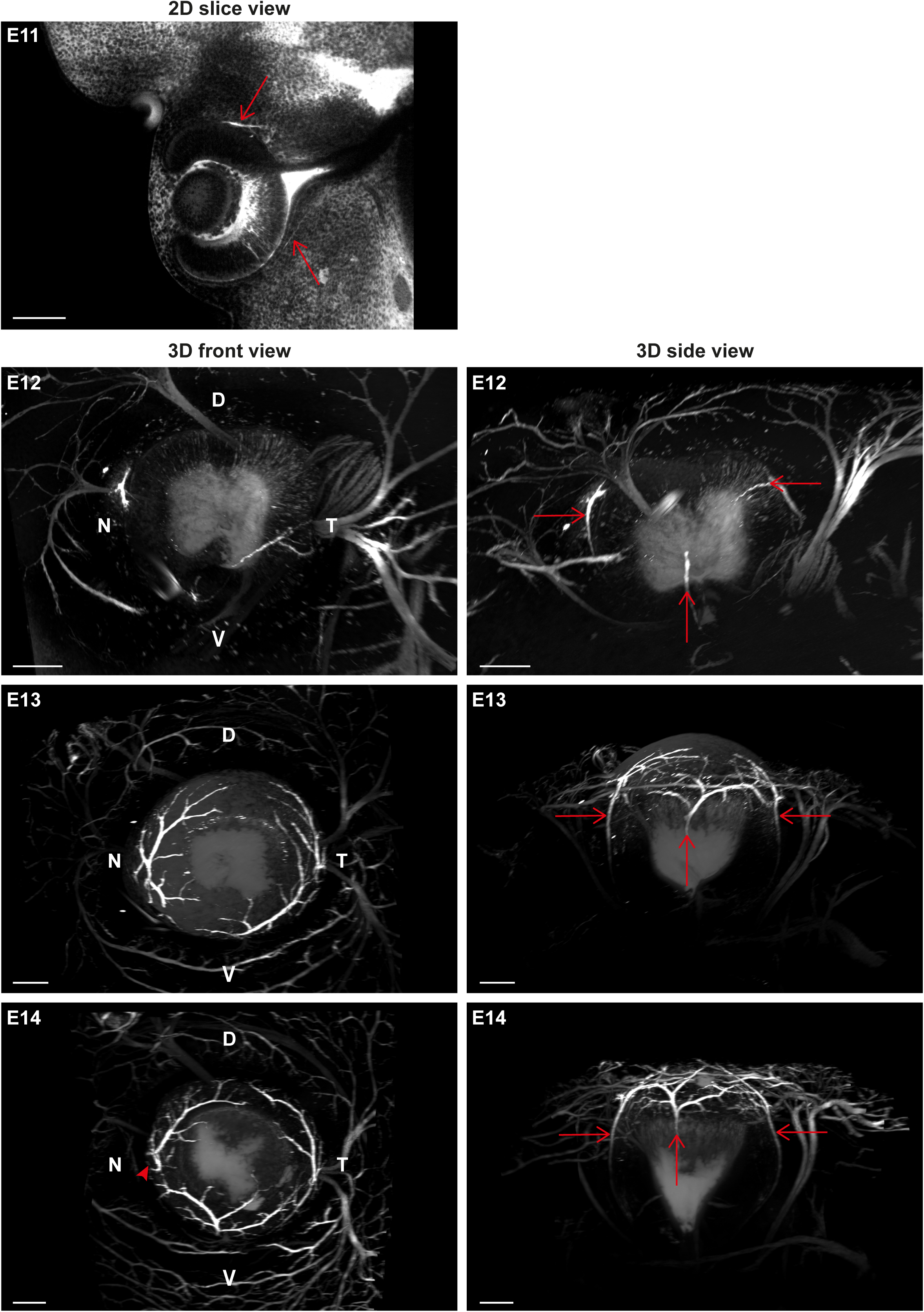
Development of innervation in mouse embryos. All embryos were cleared, and innervation was labeled using βIII tubulin antibody. Red arrows indicate the position of fiber bundles reaching the cornea. The image at stage E11 represents an optical section. The other images are 3D projections, with a coronal view on the left and a sagittal view of the eye on the right. The red arrowhead at E14 indicate the repulsion of the nasal branch by the ventral one. D: dorsal, T: temporal, N: nasal, V: ventral. Scale bar: 200 µm.

### Corneal sensory innervation has a slow paced maturation

The corneal innervation comprises various fiber types characterized by distinct functionalities, electrophysiological properties, and specific molecular markers. To investigate the developmental dynamics of these fibers, we conducted volumetric analyses from birth (P0) through adulthood (12 weeks of age, 12W), as previously described^29^. Specifically, we focused our analysis on fibers expressing *L1CAM*, *NF200*, *SP*, and *TRPM8* markers.

Initially, we visualized the morphogenesis dynamics of corneal innervation using βIII-tubulin immunolabeling, which labels all nerve fibers (**Figure 2**). At birth (P0), βIII-tubulin labeling revealed sparse and uniformly distributed innervation. While no significant changes occurred by the eyelid opening stage (P14), we observed an approximately tenfold increase in innervation volume between P14 and 4 weeks of age. Between 4 and 12 weeks, a distinct subbasal nerve plexus organization emerged, characterized by a central vortex pattern persisting into adulthood.

**Figure 2.**
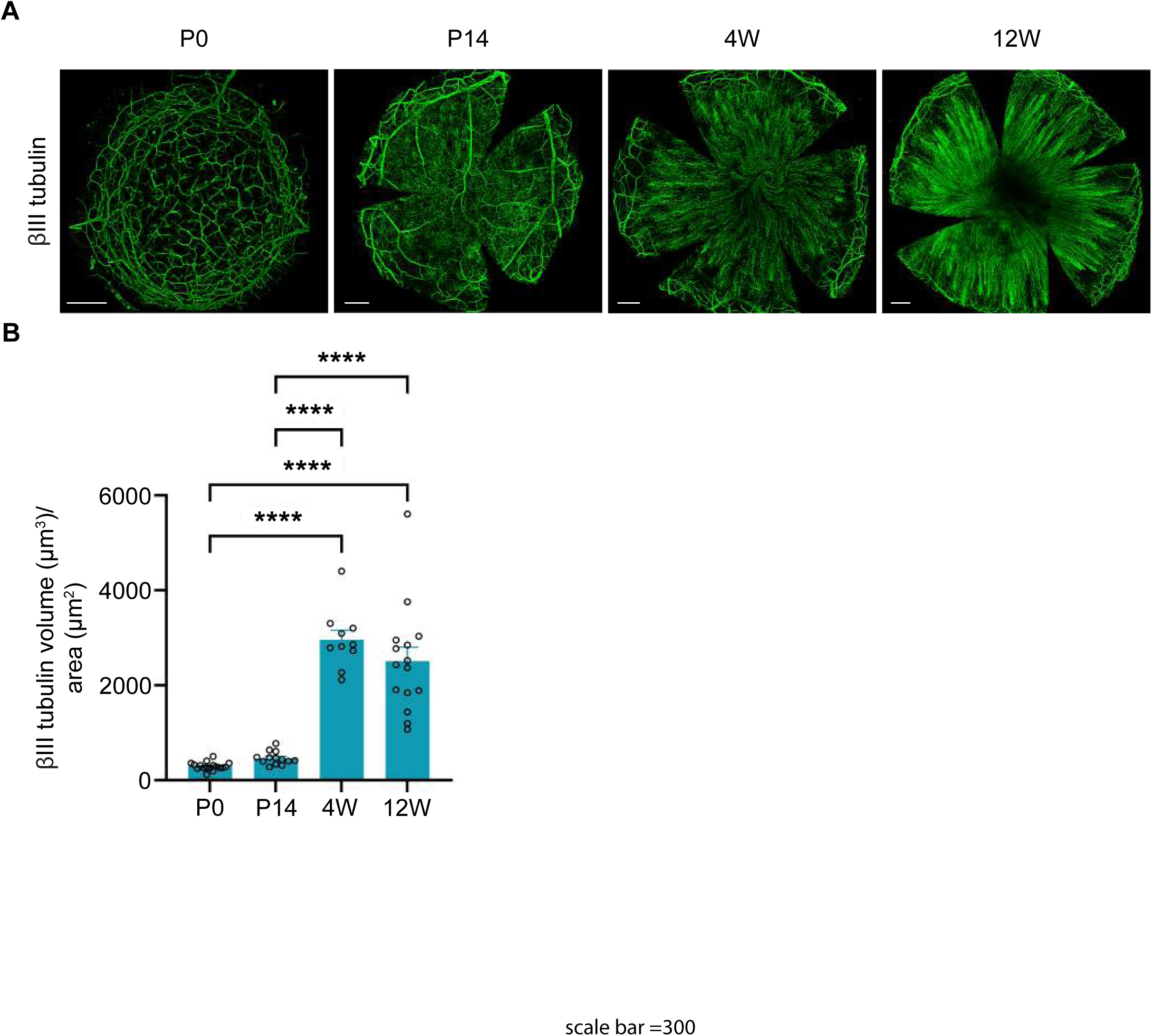
Visualization and volume analysis of the βIII-tubulin+ innervation. (**A**) The visualization of βIII-tubulin+ axons showed the progressive innervation pattern formation from P0 to 12 weeks of age. The fiber volumes are then measured for each stage using Imaris software (n=10-20). Volumes of βIII-tubulin+ fibers is drastically increased from P14 to 4 weeks of age. Data are represented as mean ±SEM. Statistical analysis using Mann Whitney test, and *p<0.05. Scale bar = 300µm.

We subsequently applied the same visualization strategy to fibers labeled with L1CAM, NF200, SP, and TRPM8. *L1CAM*+ fibers displayed patterns similar to βIII-tubulin-labeled fibers (**Figure 3**). Conversely, *NF200*+, indicative of myelinated fibers, progressively diminished during development (**Figure 4**), consistent with previous reports of the absence of myelinated fibers in mature corneas^30^. Interestingly, *SP*+ (**Figure 5**) and *TRPM8*+ (**Figure 6**) fibers were abundant at P0 and included small fibers not labeled by respectively βIII-tubulin or L1CAM.

**Figure 3.**
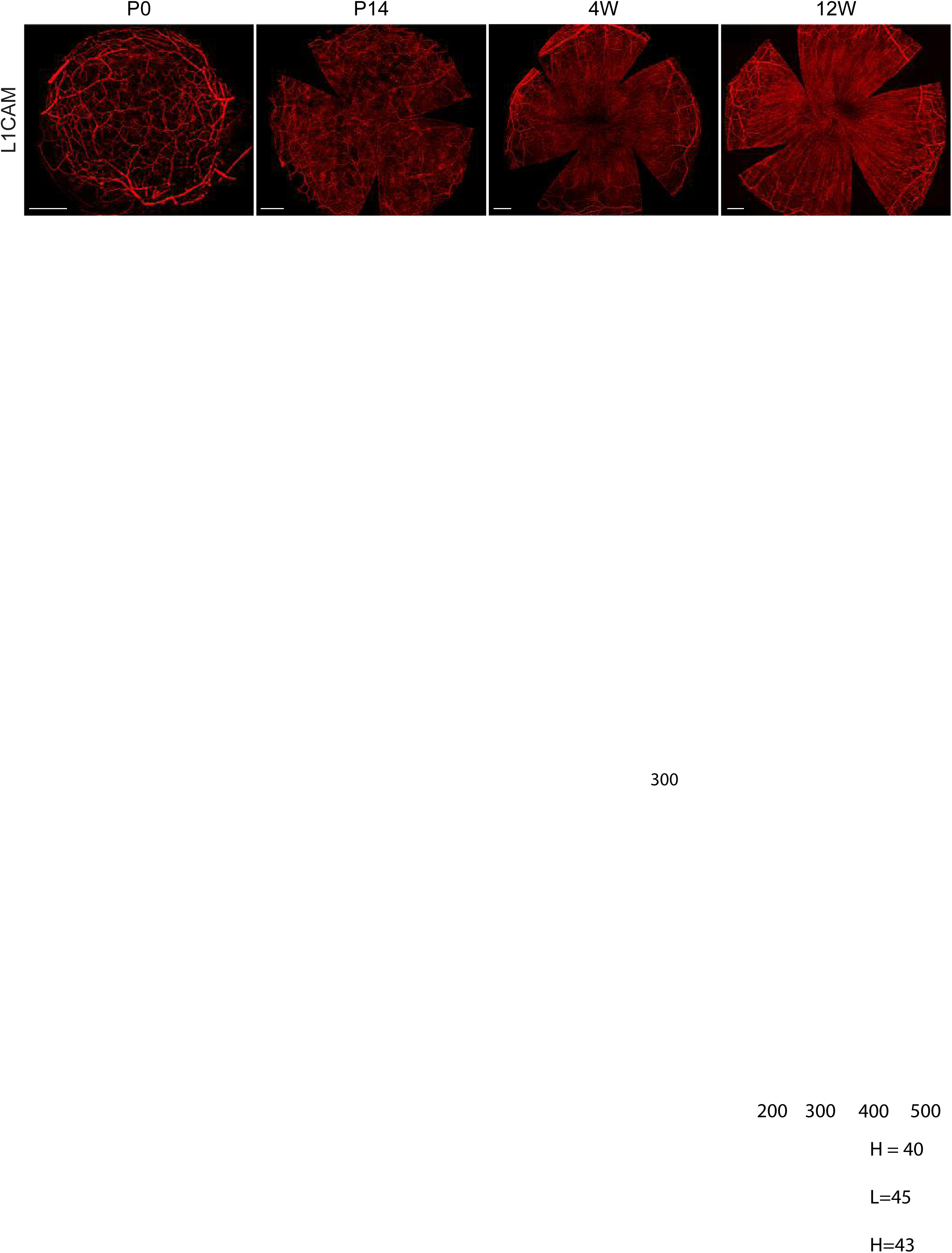
Visualization of the L1CAM+ innervation. The visualization of L1CAM+ innervation showed the progressive innervation pattern formation from P0 to 12 weeks of age. Scale bar = 300µm.

**Figure 4.**
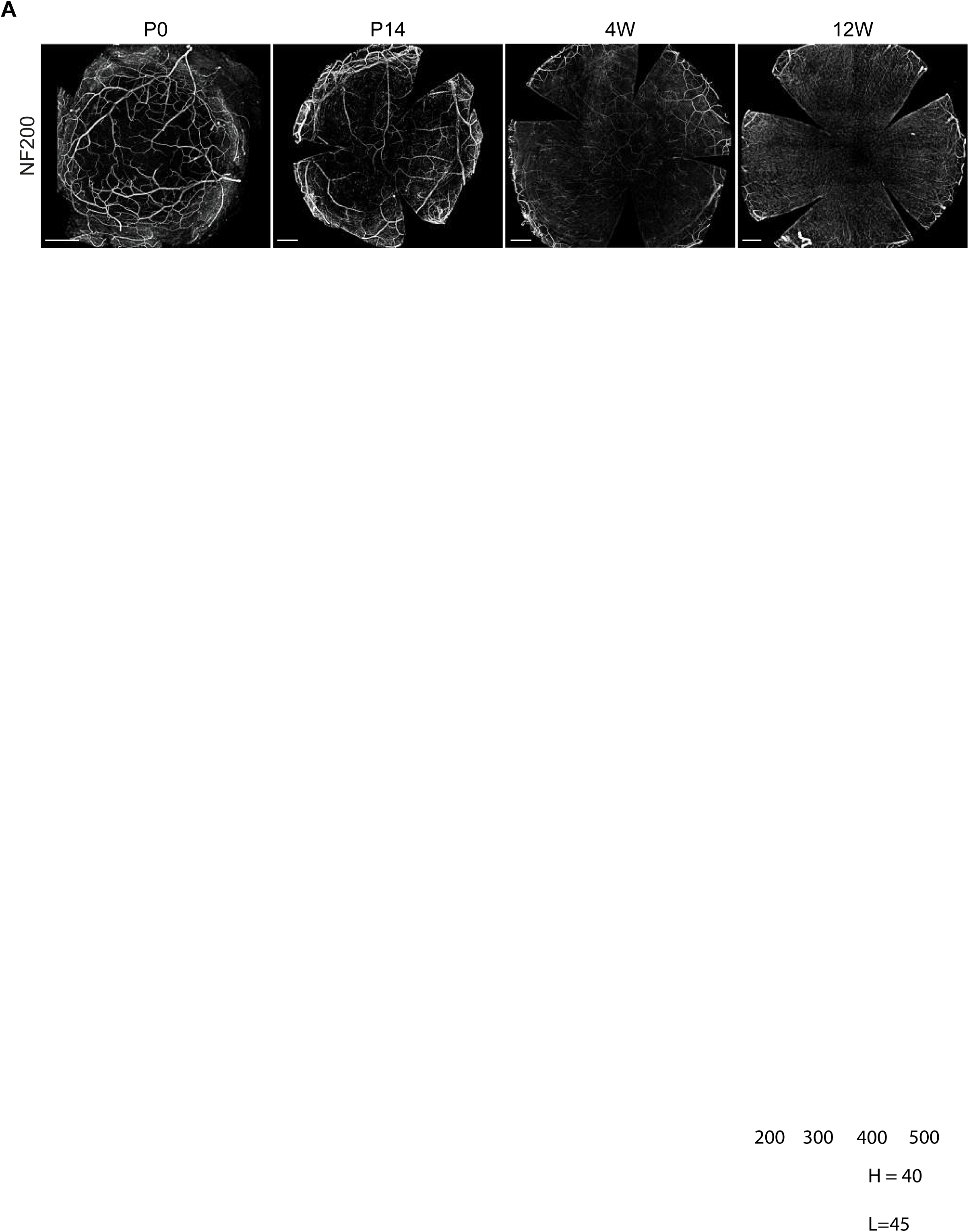
Visualization of the NF200+ innervation. The visualization of NF200+ innervation showed the progressive innervation pattern formation from P0 to 12 weeks of age. Scale bar = 300µm.

**Figure 5.**
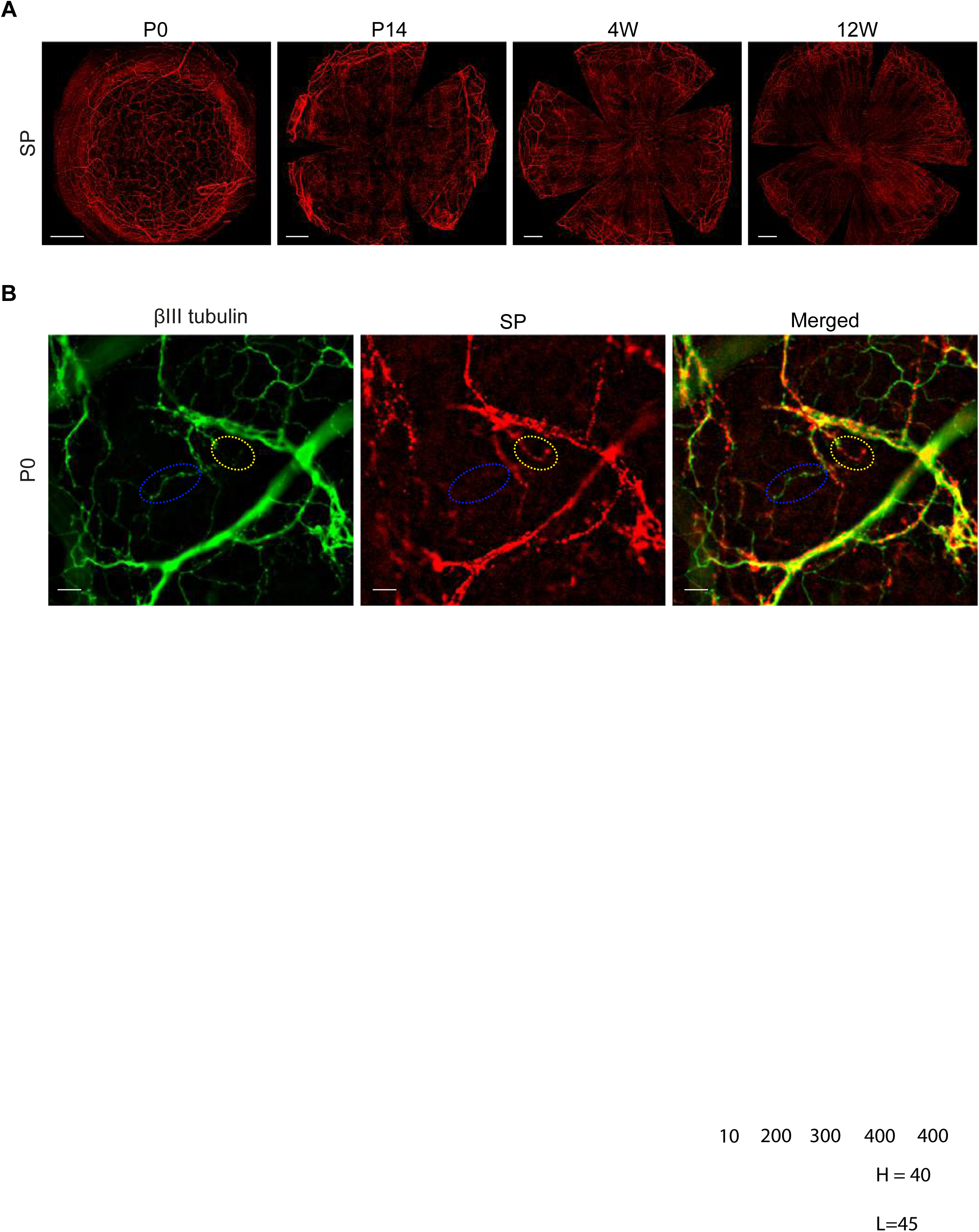
Visualization of the SP+ innervation. (**A**) The visualization of SP+ innervation showed the progressive innervation pattern formation from P0 to 12 weeks of age. (**B**) The close-up on some small terminal fibers at P0 showed the absence of βIII-tubulin in SP+ domains. Scale bar = 300µm (**A**), and 300µm (**B**).

**Figure 6.**
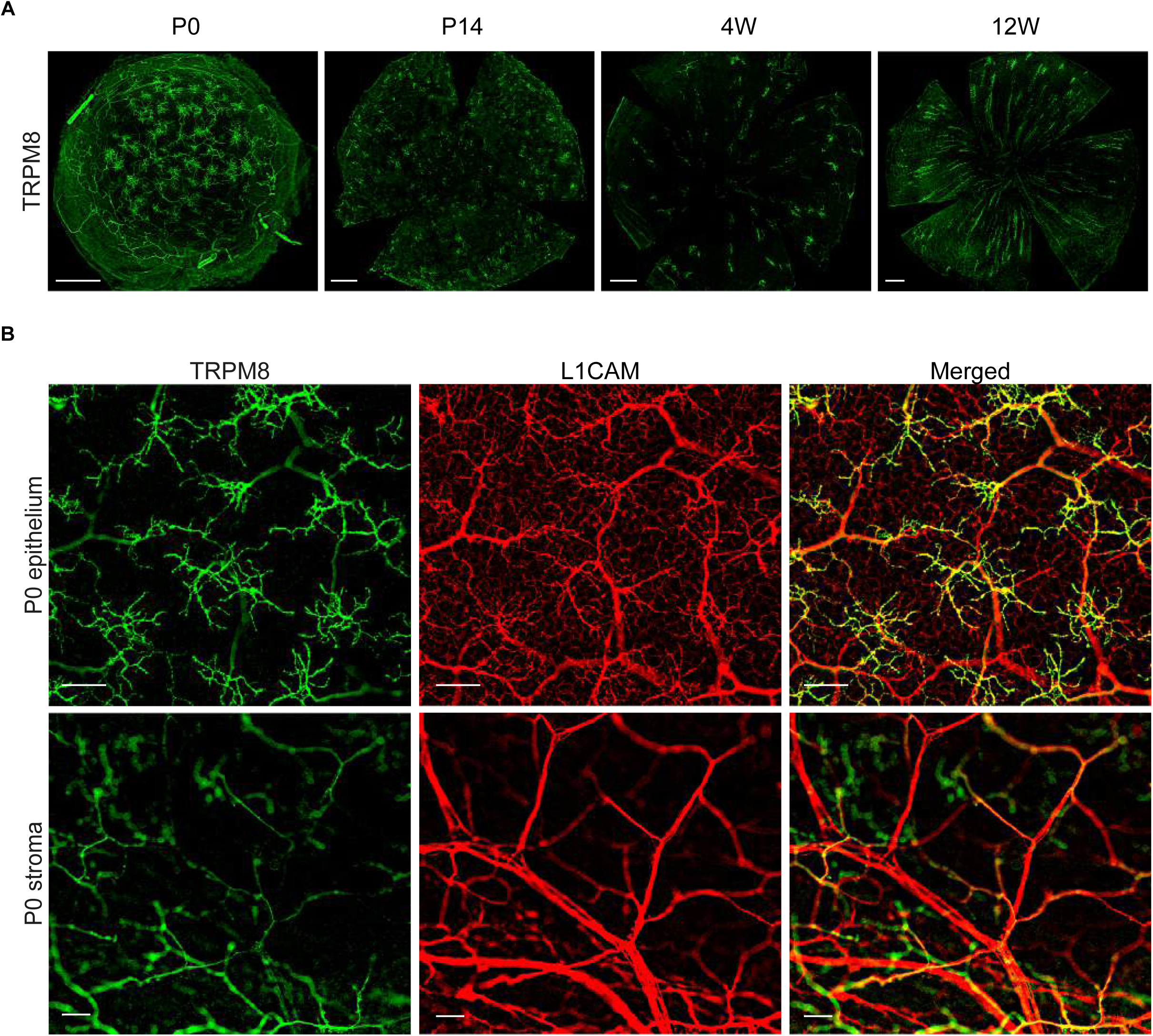
Visualization of the TRPM8+ innervation. (**A**) The visualization of TRPM8+ innervation showed the progressive innervation pattern formation from P0 to 12 weeks of age. (**B**) The close-up on some small terminal fibers at P0 in the epithelium as well as in the stroma showed the absence of L1CAM in SP+ domains. Scale bar = 300µm (**A**), and 300µm (**B**).

Quantitative volume analysis revealed fiber-specific developmental dynamics when compared to βIII-tubulin labeling (**Figure 7**). *L1CAM*+ fiber volumes peaked around eyelid opening (P14), coinciding with the initiation of corneal stratification, suggesting a role in innervation densification. *NF200*+ fibers decreased steadily over time, as anticipated. Notably, *SP*+ and *TRPM8*+ fibers initially decreased between P0 and 4 weeks, followed by a significant increase in adulthood, suggesting a refined specialization of corneal innervation during later developmental stages. Further comparative volume analyses at each developmental stage underscored these patterns (**Figure 8**).

**Figure 7.**
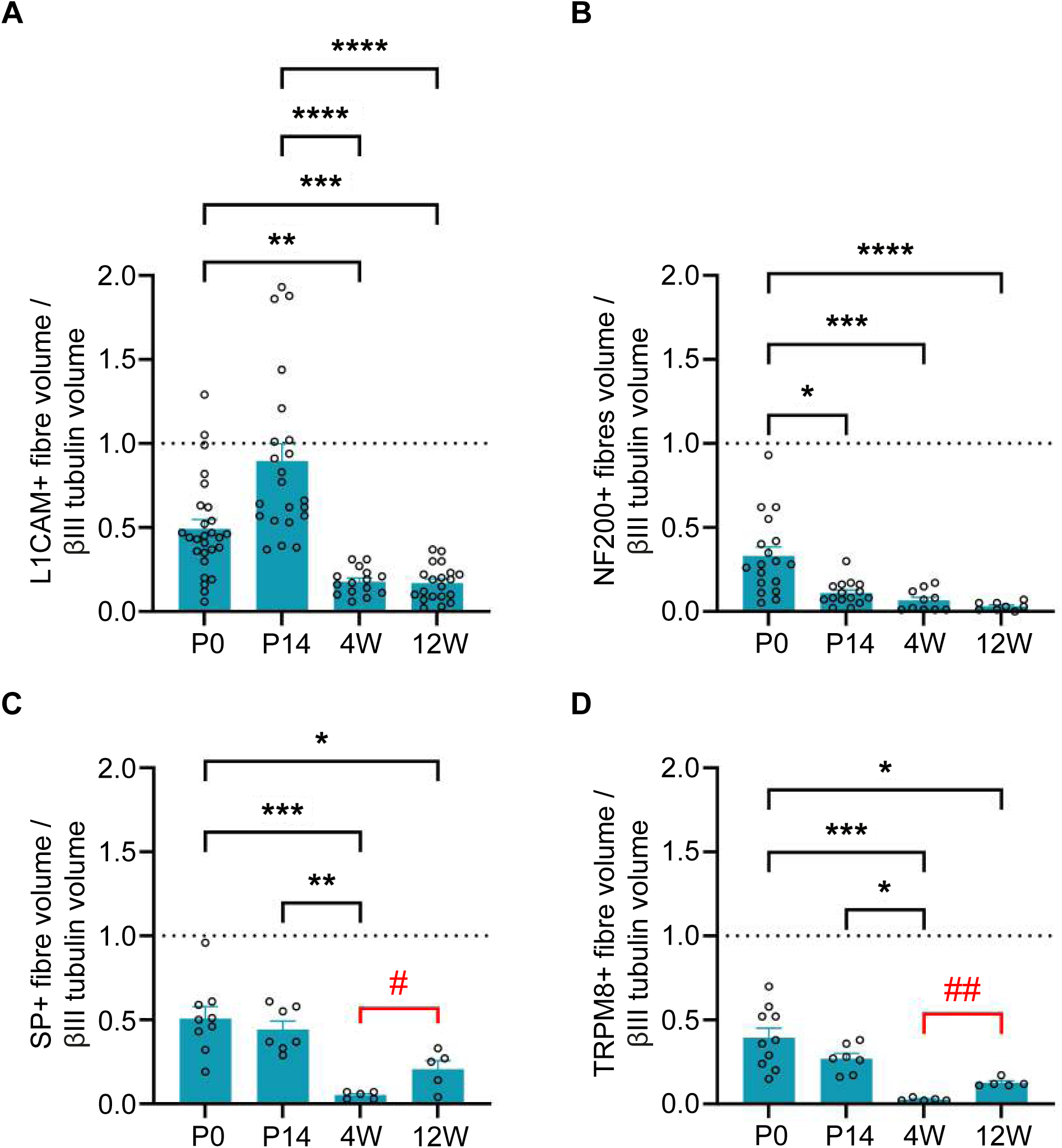
Dynamics of specific fiber types volume from P0 to 12 weeks of age. (**A**) The measurement of corneal innervation volume for each fiber types during innervation morphogenesis highlights a large increase of L1CAM+ volume at P14, then a decreased to a stable level. (**B**) NF200+ volume decreases steadily during the morphogenesis process. (**C**) SP+ decreases from P14 to 4 weeks of age, but significantly increases by 12 weeks. (**D**) TRPM8+ volume follows the same profile than the SP+ one (n=10-20). Data are represented as mean ±SEM. Statistical anlaysis using two-way ANOVA with uncorrected Fisher’s, and *p<0.05; ***p<0.001.

**Figure 8.**
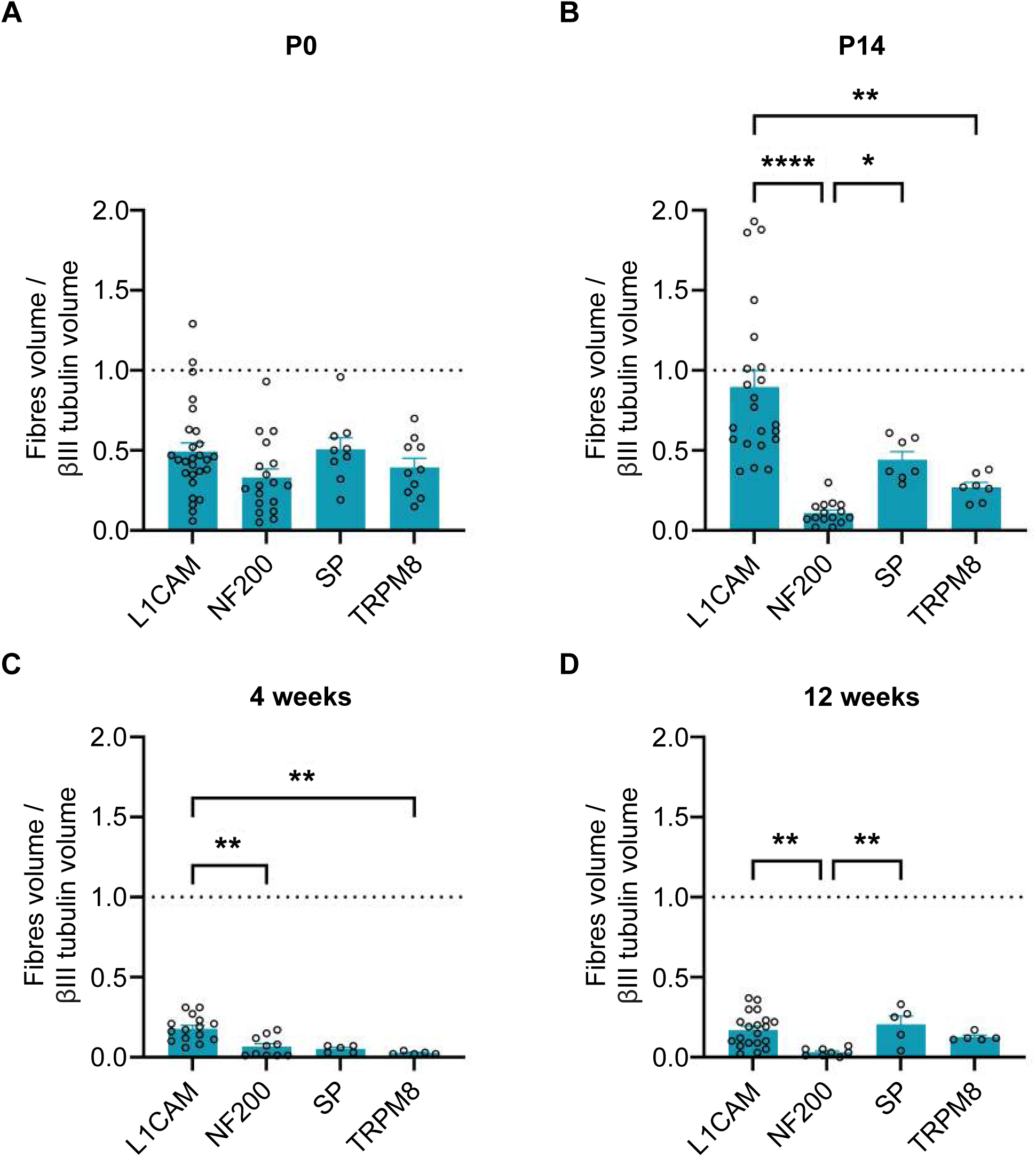
Dynamics of specific fiber types volume from P0 to 12 weeks of age. (**A**) The visualization by stages is convenient to interrogate the proportion of each fiber type at any given timepoint. At P0 each marker represented the same proportion of the innervation. (**B**) At P14, L1CAM+ volume was close to βIII-tubulin, showing a strong expression of this marker. (**C**) At 4 weeks, L1CAM was still the most expressed marker, but NF200, SP and TRPM8 were very lowly expressed. (**D**) At 12 weeks, while NF200 was close to not detected, L1CAM, SP and TRPM8 represented a similar volume (n=10-20). Data are represented as mean ±SEM. Statistical anlaysis using two-way ANOVA with uncorrected Fisher’s, and *p<0.05; ***p<0.001.

Additionally, we examined corneal sensory sensitivity across development (**Figure 9**). Sensitivity increased progressively, reaching a plateau around 3 months of age, with minor incremental changes continuing until 8 months.

**Figure 9.**
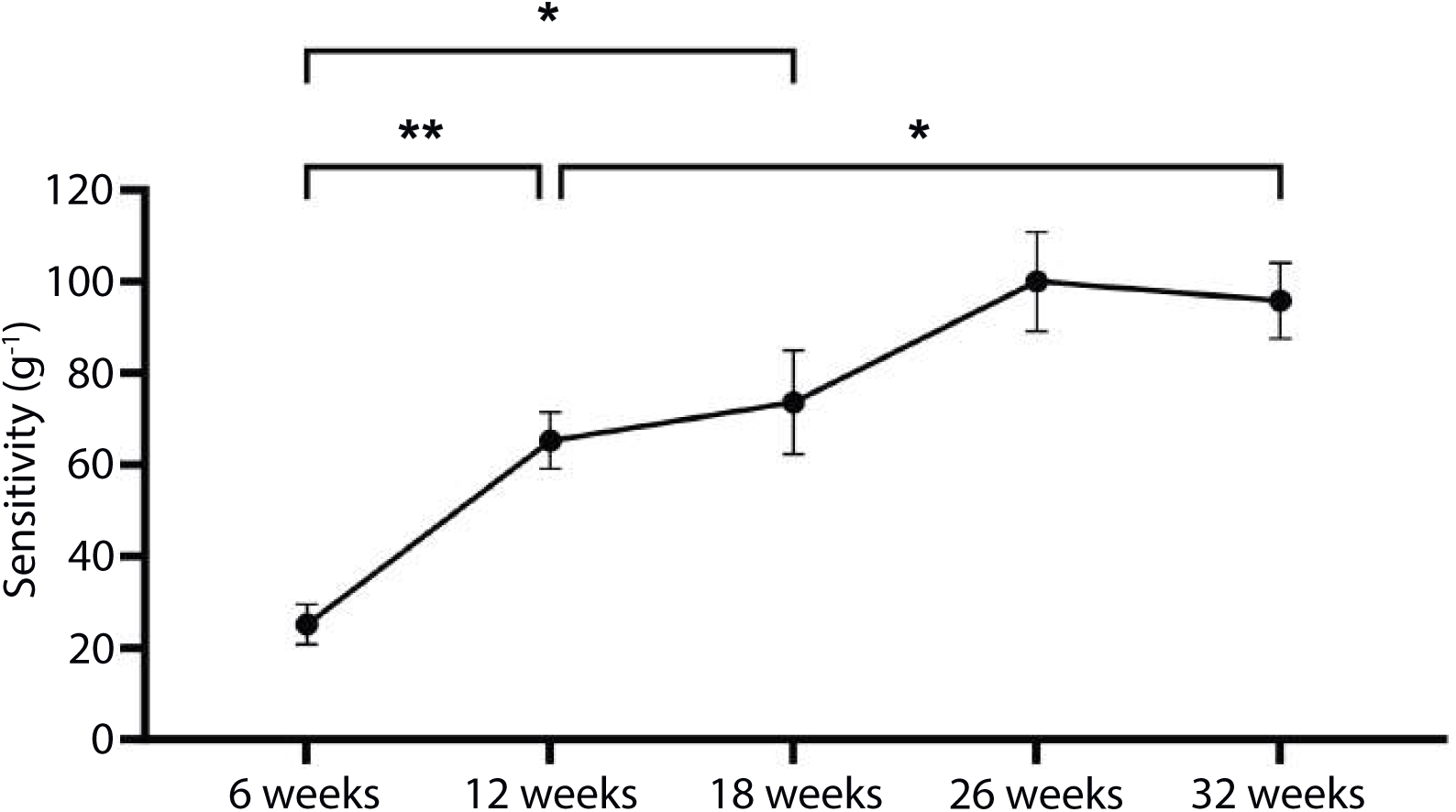
Corneal sensitivity evolution from 6 to 32 weeks of age. Central corneal sensitivity is monitored up with Von Frey filaments. The sensitivity increases gradually from 6 to 18 weeks of age. Then, the sensitivity reaches a plateau by 26 weeks of age (n=13-15). Data are represented as mean ±SEM. Statistical analysis using mixed-effect analysis with Sidak’s multiple comparisons test. *p<0.05 and ***p<0.001.

Together, our findings illustrate that corneal innervation undergoes slow morphogenesis and maturation, with pronounced developmental dynamics between 4 and 12 weeks of age.

To understand the impact of corneal innervation loss, and its physiological regeneration, we used two clinically-relevant models, the axotomy and the epithelial abrasion.

### Characterization of the murine axotomy model recapitulating penetrating keratoplasty

During corneal transplantation procedures, called penetrating keratoplasty, corneal innervation is inevitably severed around the transplanted region, necessitating subsequent nerve regeneration^31^. To understand the effects of this innervation loss on corneal epithelial cells and to characterize the regenerative process, we employed our previously established axotomy model^32^ (**Figure 10**, **Figure 11**).

**Figure 10.**
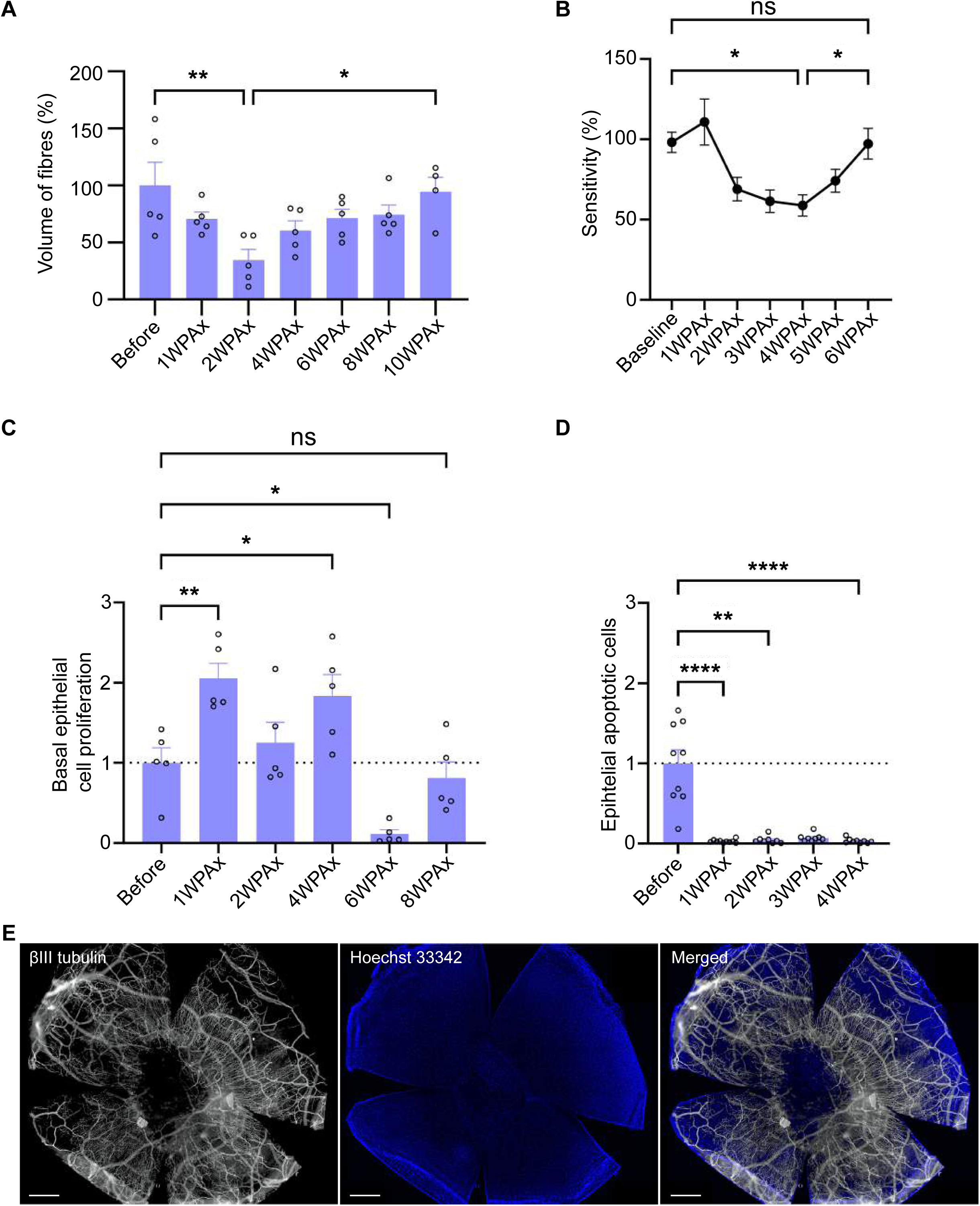
Characterization of the corneal parameters after axotomy. Fiber volume (**A**), corneal sensitivity (**B**), and the number of proliferating (**C**) and apoptotic (**D**) cells in the basal epithelium were measured at various time points after axotomy. Example of a cornea exhibiting an ulceration in the epithelium 2 weeks after surgery. WPAx: weeks post-axotomy. Scale bar: 400 µm. *p<0.05, **p<0.01, and ****p<0.0001.

**Figure 11.**
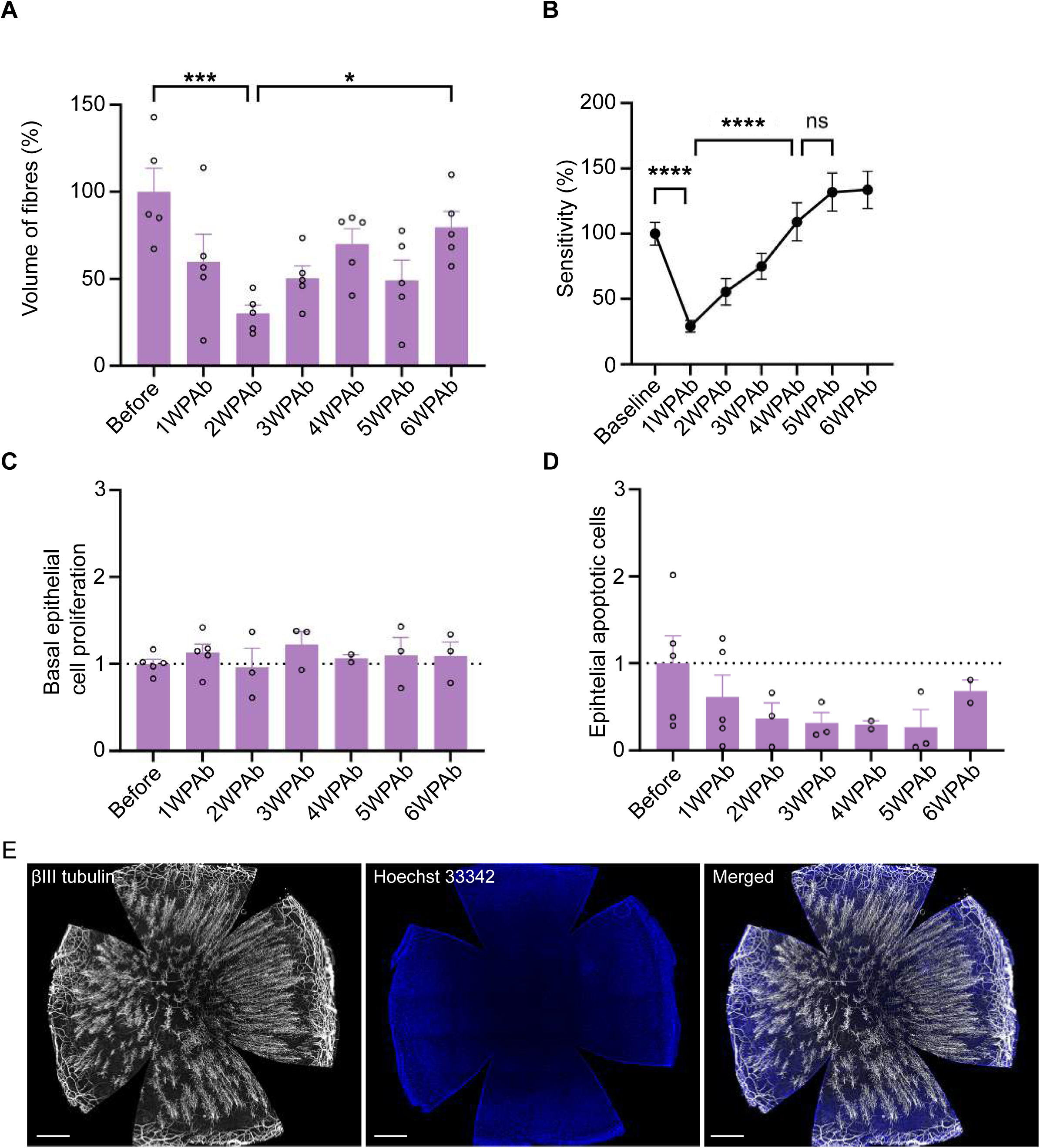
Characterization of the cornea and its innervation after abrasion. Fiber volume (**A**), corneal sensitivity (**B**), and the number of proliferating (**C**) and apoptotic (**D**) cells in the basal epithelium were measured at various time points after abrasion. Image of a cornea 4 weeks after abrasion. WPAb: weeks post-abrasion. Scale bar: 500 µm. *p<0.05, **p<0.01, ***p<0.001, and ****p<0.0001.

We compared the innervation volume before axotomy with subsequent regeneration phases to assess the regenerative dynamics and their impact on epithelial cells. Our analysis demonstrated a significant reduction in innervation volume immediately after axotomy. Innervation volume gradually recovered, achieving pre-injury levels approximately 10 weeks post-axotomy (10WPAx) (**Figure 10A**). Interestingly, while structural nerve regeneration spanned approximately 10 weeks, functional recovery, as indicated by sensory sensitivity restoration, occurred sooner (**Figure 10B**). Sensory functionality was restored by 6 weeks post-axotomy, thus preceding the full structural regeneration by 4 weeks. This finding highlights a notable disconnect between anatomical innervation volume and sensory function.

To explore the impact of nerve injury on corneal epithelial cell homeostasis, we measured epithelial cell proliferation and apoptosis. Our analysis revealed substantial dysregulation in epithelial cell proliferation, which significantly increased during the first four weeks post-axotomy (**Figure 10C**). In parallel, apoptosis was markedly suppressed over this period, causing considerable disruption to normal epithelial homeostasis (**Figure 10D**). Occasionally, this disruption resulted in severe epithelial damage, including ulceration (**Figure 10E**). However, such extreme cases were infrequent and thus excluded from the quantitative analysis.

Collectively, our findings highlight a dissociation between structural nerve regeneration and functional recovery, as well as significant nerve injury-induced disturbances in epithelial cell proliferation and apoptosis dynamics.

### Characterization of the murine abrasion model recapitulating photorefractive keratitis model

To further characterize corneal nerve regeneration, we utilized a clinically relevant mouse model mimicking photorefractive keratitis, achieved by mechanical epithelial abrasion^32^. Our analyses began one week post-abrasion to ensure complete epithelial healing^33^, and extended over subsequent weeks.

Following epithelial abrasion, we observed a marked reduction in corneal innervation volume, reaching its lowest at two weeks post-abrasion (2WPAb) (**Figure 11A**). Interestingly, corneal sensitivity was lowest one week post-abrasion (1WPAb), preceding the lowest structural innervation volume, recorded at 2 weeks post-abrasion (2WPAb) (**Figure 11B**). Sensitivity fully recovered within the subsequent three weeks, highlighting a temporal mismatch between functional sensitivity recovery and structural nerve regeneration.

Additionally, we assessed corneal epithelial homeostasis throughout nerve regeneration. Phospho-histone3+ basal epithelial cell counts remained stable during the regeneration process (**Figure 11C**). Although a trend toward reduced apoptosis was noted, statistical analysis using ANOVA revealed no significant differences across the examined time points (**Figure 11D**). These findings provide critical insights into the distinct timelines of structural and functional corneal nerve recovery and highlight subtle shifts in epithelial homeostasis during regenerative processes post-injury.

The structural innervation gradually regenerated over five weeks following abrasion. Histological evaluations at four weeks post-abrasion revealed incomplete nerve regeneration within the central cornea, characterized by nerve-free areas interspersed with dense nerve fiber clusters (**Figure 11E**).

### Distinct regenerative process after abrasion or axotomy

As the axonal regenerative process appeared specific to the type of corneal innervation injury sustained, we examined the expression of 12 neural genes commonly associated with neurogenesis: *Atf3*, *Calca*, *L1cam*, *Nf200*, *Ngfr*, *Nrp1*, *Ntrk1*, *Piezo2*, *Robo1*, *SP*, *Trpm8*, and *Tubb3*. Despite the relatively small population of trigeminal ganglion neurons innervating the cornea, we selected this tissue to evaluate gene expression (**Figure 12**).

**Figure 12.**
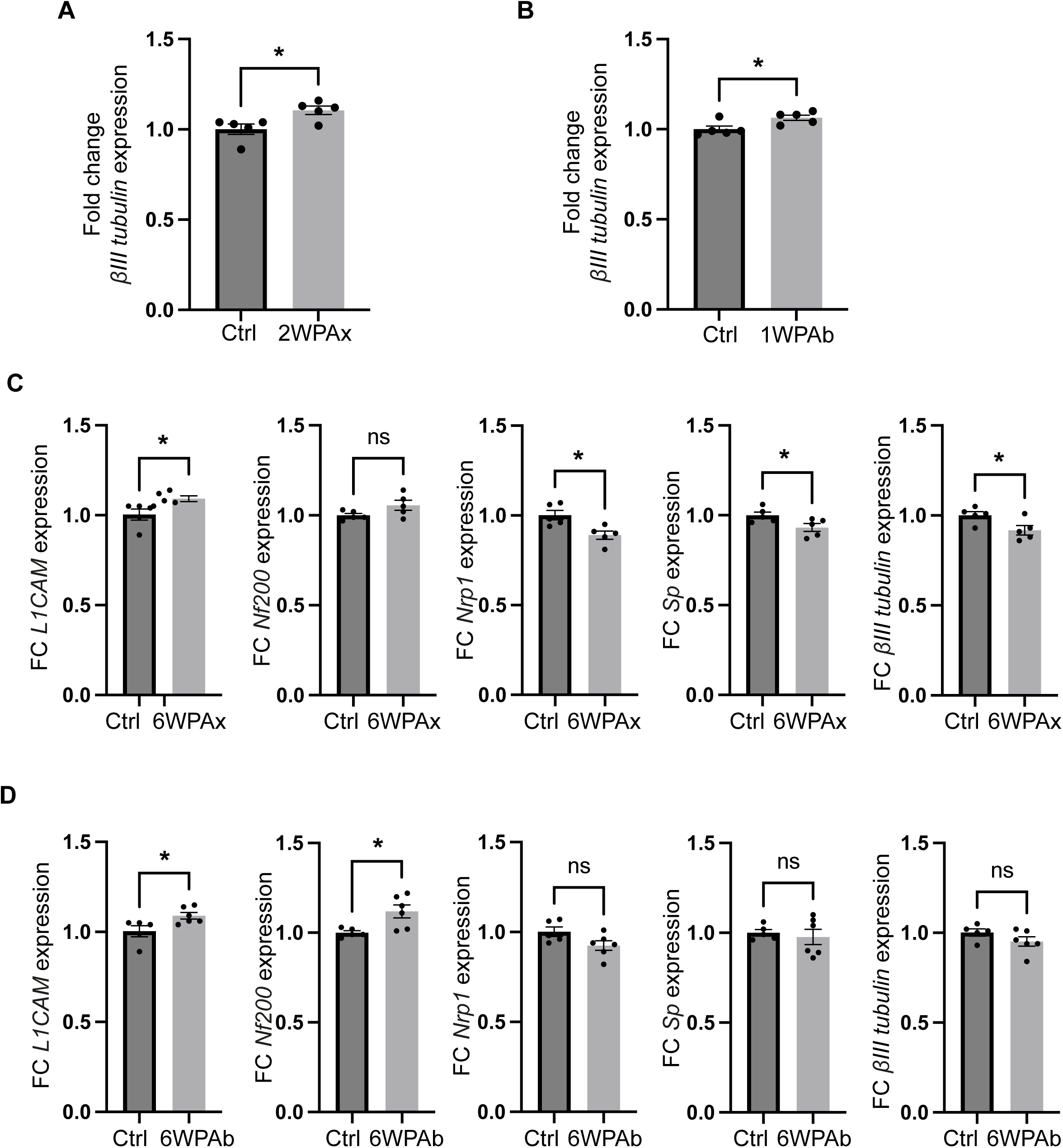
Neural gene expression is modulated by innervation injury. (**A**) *βIII tubulin* expression is over expressed 2 weeks post axotomy (2WPAx), similarly to 1 week post abrasion (1WPAb) (**B**). (C) *L1cam*, *Nrp1*, *Sp* and *βIII tubulin* expression are modulated 6WPAx, but not *Nf200*. (**D**) *L1cam* and *Nf200* expression are modulated 6WPAb, but not *Nrp1*, *Sp*, or *βIII tubulin*. Data are represented as mean ±SEM. Statistical analysis using t-test.ns, non significant, *p<0.05.

Initially, we quantified βIII tubulin expression at time points corresponding to reduced corneal sensitivity, i.e. two weeks post-axotomy (**Figure 12A**) and one week post-abrasion (**Figure 12B**). Both injury types resulted in increased βIII tubulin expression. Considering that the 6-week post-injury interval seemed critical for active innervation regeneration, we assessed the expression of the 12 selected genes at this specific time point. Among these, seven genes showed no significant differences compared to control conditions. Four genes exhibited altered expression levels following axotomy (**Figure 12C**), while only two genes showed modulation after abrasion (**Figure 12D**). Notably, L1cam was the only gene consistently upregulated after both injury types, highlighting the specificity of regenerative processes in response to distinct corneal injuries.

## Discussion

The integrity and functionality of the cornea critically depend on its dense sensory innervation. Beyond facilitating sensation, corneal innervation plays an essential role in maintaining corneal transparency, structural integrity, and promoting efficient wound healing. Damage to or loss of these nerve fibers can lead to severe clinical conditions, notably neurotrophic keratitis, characterized by compromised wound healing, progressive corneal degeneration, ulceration, and potential vision impairment or loss. Currently, a limited understanding of underlying mechanisms restricts the development of innovative treatments. This report establishes foundational knowledge essential for comprehending the regeneration processes of corneal innervation.

Corneal nerve morphogenesis and maturation span from embryonic day 12 (E12) through 12 weeks postnatally. Our findings reveal that trigeminal nerve fibers first arrive at the cornea by E12, initially exhibiting delayed innervation in the dorsal region. By E14, however, the entire corneal periphery is fully innervated. Notably, while innervation density increases progressively until eyelid opening, fiber-specific differentiation and functional maturation predominantly occur post-eyelid opening. Concurrent changes in fiber-type distribution and sensory sensitivity from 2 to 12 weeks underscore the gradual maturation of corneal sensory innervation.

Marker-specific analyses demonstrated early widespread expression of L1CAM, a molecule associated with axon guidance and adhesion, with significant decreases observed by 4 and 12 weeks as fibers became thinner and less intensely labeled. Conversely, NF200, indicative of myelination, substantially declined postnatally, becoming nearly absent by adulthood. SP, involved in nociception and tear secretion, markedly decreased post-eyelid opening but exhibited a slight rebound toward adulthood. TRPM8-positive fibers, sensitive to cold and osmotic stimuli, followed a similar expression trend—initial reduction after eyelid opening, followed by modest increases during adulthood, predominantly localized within epithelial nerve terminals. Analysis of fiber type proportions revealed a uniform distribution at birth, with NF200+ fibers significantly diminishing from P14 onward. Initially dominant L1CAM+ fibers decreased proportionally between 4 and 12 weeks. Simultaneously, SP+ and TRPM8+ fibers proportionally increased, eventually reaching equal representation with L1CAM+ fibers by adulthood, indicating progressive specialization and refinement of corneal innervation during maturation. Together, our results demonstrate the importance of L1CAM during innervation morphogenesis.

Interestingly, the dynamics of sensitivity loss and epithelial cell homeostasis disruption following corneal innervation injury appear dependent on the type of nerve damage inflicted. Axotomy, which triggers Wallerian degeneration^32^, leads to a prolonged and significant impact on epithelial apoptosis. In contrast, epithelial abrasion, affecting only the terminal ends of nerve fibers, has a comparatively mild effect on apoptosis. Moreover, corneal ulceration was exclusively observed following axotomy. These findings underscore the variable consequences resulting from different modalities of innervation loss, indicating distinct underlying pathological mechanisms. Consequently, the molecular environments following axotomy and abrasion differ significantly. We hypothesize that epithelial abrasion, known to promote epithelial plasticity^34^, likely influences the pathological environment substantially, potentially modulating the overall regenerative and healing response.

Our findings on neural gene expression following injury further support this hypothesis. Specifically, epithelial injury appears to be associated with decreased expression of Nrp1, a factor known to play a crucial role in the development of stromal and epithelial innervation^35^. Importantly, abrasion injuries are confined to the epithelium, unlike axotomy, which affects deeper neural structures.

Additionally, the increased expression of L1cam and Nf200 observed after abrasion resembles a recapitulation of embryonic states when innervation initially forms. It is conceivable that epithelial injury triggers epithelial plasticity, leading to a more naïve, embryonic-like state of innervation. This naïve innervation may then undergo differentiation, ultimately restoring full functional capacity.

Collectively, these observations highlight that the corneal sensitivity and nerve fiber volume recovery post-injury are not strictly correlated. Instead, nerve fiber identity and phenotype plasticity likely influence corneal sensitivity restoration. Thus, our mouse models provide valuable insights into nerve fiber regeneration dynamics, offering potential therapeutic targets for clinical conditions such as neurotrophic keratopathy and dry eye disease.

## Material and Methods

### Animals included in this study

All the experiments conducted on mice were approved by the local ethical committee and the Ministère de la Recherche et de l’enseignement Supérieur (authorization 2016080510211993 version2). All of the procedures were carried out in accordance with the French regulation for the animal procedure (French decree 2013-118) and with specific European Union guidelines for the protection of animal welfare (Directive 2010/63/EU). Mice were housed in plastic boxes, on a standard light cycle (12h light, 12h dark), with food and water ad libitum, in a 40-60% relative humidity environment and a 21°C-22°C ambient temperature. Embryos were collected on ice between E11 and E14. Swiss/CD1 female mice (RjOrl:SWISS, Janvier Labs, France) were used for all experiments. Mice were euthanazied by cervical disclocation. Puppies were ethanazied by beheading.

### Embryo treatment

Embryos were fixed in a fresh solution of 4% paraformaldehyde (PFA) at 4°C overnight and rinsed in phosphate-buffered saline (PBS). The embryos were then incubated in a blocking solution composed of 5% goat serum goat serum (GS) (Merck, S26-100mL), 2.5% fish skin gelatin (FSG) (Sigma-Aldrich, #G7765) and 0.5% Triton X-100 in PBS for 24h at room temperature. Embryos were then incubated with the primary antibody (antibody anti-βIII tubulin) for 7 days at 4°C. Embryos were rinsed and incubated with the secondary antibody for 7 days at 4°C. After rinsing, they were transparentized using CUBIC reagent 1 (R1) clearing solution (refer to Susaki et al. 2015). First, the embryos were incubated in a R1 ½ solution (diluted with dH2O) at 37°C. After two hours, the solution was replaced by R1 and the embryos kept at 37°C. The solution was changed every day for 5 to 7 days until the embryos were transparent.

### Embryo imaging

Embryos were imaged using a Lightsheet Z1 (Zeiss) with a 5X objective. Embryos were suspended upside down using a homemade hook, attached to a holder supplied with the microscope. The embryos were immersed in R1 solution in the microscope tank. The images were acquired with a 1.06 optical zoom.

### Sensitivity

Sensitivity of the central cornea was assessed *in vivo* using Von Frey filaments (Bioseb, France, #vio-VF-M) as previously described^36^. Succinctly, the nylon filaments of increasing strength were applied until the mouse blinked.

### Tissue collection and processing

For each sample, two trigeminal ganglions from the wounded side of two mice were collected and pooled in lysis solution, snap frozen then stored at −80°C. Eyes were collected using curved scissors to cut the optic nerve. They were then fixed in 4% PFA (Antigenfix) for 20 minutes, then washed in PBS. The eyes were then dehydrated for 2H in 50% ethanol/PBS and stored in 70% ethanol/PBS at 4°C.

### Corneal abrasion

Corneal abrasions were performed as previously described^33^. Mice were anesthetized by intraperitoneal injection with a mix of ketamine (80mg/kg, Imalgene® 1000, Centravet) and medetomidine (1mg/kg, Domitor®, Centravet). The center of the corneal epithelium of the righteye was circulary removed using an ocular burr (Algerbrush II, reference BR2-5 0.5 mm, The Alger Company, USA). Immediatly after, the abrasion was checked using a fluorescein solution (1% in PBS, Sigma-Aldrich) under cobalt blue light. After abrasion, a drop of Ocrygel (TVM) was applied on both eyes, mice were treated with an analgesic solution of buprenorphine (0.1mg/kg, Burprecare®, Centravet). The wellbeing of the animals was monitored in the following days.

### Corneal nerves axotomy

Axotomy of the corneal nerves were performed as previously described^32^. Mice were anesthetized by intraperitoneal injection with a mix of ketamine (80mg/kg, Imalgene® 1000, Centravet) and medetomidine (1mg/kg, Domitor®, Centravet). Under a binocular loupe, lift the eye out of the orbit using smooth, curved forceps. Next, a 2.5 mm biopsy punch is applied vertically to the cornea with gentle pressure and twisted several times. If the eye ruptures, the animal must be euthanized. Once the punch is removed, a visible circle should be seen on the cornea. Ocular gel is applied to both eyes while the mouse recovers on a heated pad at 37°C. The animal is monitored for the next two days.

### RT-qPCR on trigeminal ganglion

Samples composed of two pooled trigeminal ganglions were used for the RNA extraction. The quantitative PCR experiments were performed using a TaqMan detection protocol on the QuantStudio 12K Flex Real-Time PCR System (Life Technologies). The selected TaqMan assays (ThermoFisher Scientific) for each target and reference gene are listed in **Table 1**. For qPCR reactions, 3ng of cDNA was mixed with TaqMan FastAdvance Master Mix and the corresponding TaqMan assay in a final volume of 10 µL. Samples were then loaded onto 384-well microplates, and PCR was carried out for 40 cycles (50°C for 2 min; (95°C for 1 s, 60°C for 20 s) ×40). To verify the absence of genomic DNA contamination, negative controls lacking reverse transcriptase were included. Data normalization was performed using three reference genes (*Hprt1*, *Ppia*, and *Ubc*), with the two most stable genes (*Hprt*/*Ppia*) identified using GenEx software (MultiD). Each measurement was conducted in duplicate, and Ct values were used for analysis. The relative gene expression ratio was calculated using the ^ΔΔ^Ct method.

**Table 1.** List of genes and their identification used for the qPCR analysis.

| Gene | ID |
| --- | --- |
| **Atf3** | Mm00476033_m1 |
| **Calca** | Mm00801463_g1 |
| Hprt | Mm00446968_m1 |
| **L1cam** | Mm00493049_m1 |
| **Nf200** | Mm01191455_m1 |
| **Ngfr** | Mm00446296_m1 |
| **Nrp1** | Mm00435372_m1 |
| **Ntrk1** | Mm01219406_m1 |
| **Piezo2** | Mm01265861_m1 |
| Ppia | Mm02342430_g1 |
| **Robo1** | Mm00803879_m1 |
| **SP** | Mm01166996_m1 |
| **Trpm8** | Mm01299593_m1 |
| **Tubb3** | Mm00727586_s1 |
| Ubc | Mm01198158_m1 |

### Immunofluorescence labeling on whole cornea

Previously collected eyes were rehydrated in 50% ethanol for two hours and rinced in PBS. Corneas were dissected just above the ciliary body, along the limbal region using microdissection scissors. They were then immersed in blocking and permeabilization solution in FSG 5% (Sigma-Aldrich, reference G7765) and goat serum (GS) 5% (Thermo Fisher Scientific, reference 16,210,064) in 0,5% PBS/Triton X-100 at room temperature. Then the corneas were incubated overnight at 4°C with primary antibodies and then secondary antibodies. After washing, nuclei were stained with Hoechst for 10 minutes. The corneas were incised at four cardinal points using a 15C carbon steel surgical blade (reference 0221, Swann-Morton) and then flat-mounted in Vectashield medium (Vector Laboratories, H-1000), with the epithelium positioned against the coverslip. Apoptotic cells were labeled using the DeadEnd™ Fluorometric TUNEL System (Promega, reference G3250) on whole corneas. Briefly, freshly fixed corneas were incubated with proteinase K for 10 minutes at room temperature. They were then incubated in the rTdT reaction mix, consisting of 10% Fluorescein dUTP Nucleotide mix and 2% rTdT enzyme in Equilibration Buffer, for 60 minutes at 37°C with gentle shaking (70 rpm). To stop the reaction, samples were immersed in 2X SCC Buffer for 3 minutes at room temperature. If the TUNEL assay is combined with another marker, immunofluorescence is subsequently performed as usual.

### Images acquisition and processing

The acquisition of the images was performed as previously described^36^. A Leica Thunder Imager Tissue microscope was used to acquire the whole-cornea images, using the navigator module with the large volume computational clearing (LVCC) process. The LAS X software (version 3.7.4) was used to obtain the images using a 20X/0.55 objective. Then, to process the images Imaris Bitplane software (version 10.1) was used. All of the images from a single panel were acquired and processed with the same parameters.

### Innervation volume measurement and cell counting

A centered circle covering 50% of the corneal area was drawn using the Fiji measurement plug-in (FIJI (RRID:SCR_002285)). The corneal diameter was measured twice for each sample, and the mean value was used to determine the circle’s radius. A 50% circular crop of each cornea was then generated using the *Crop* plug-in, followed by the *Clear Outside* plug-in. The cropped images were converted to .ims format using the Imaris converter. Innervation density was quantified using the surface tool in Imaris. Once the surface was defined, volume data (expressed in μm³) were extracted for statistical analysis. Apoptotic or proliferating cells were manually counted on Fiji using the *Cell Counter* plug-in.

### Statistical analysis

Data analysis was performed using GraphPad Prism software (version 10.2.2). Results are presented as mean ± SEM. Significant p-values were represented as *p < 0.05, **p < 0.01, ***p < 0.001, ****p < 0.0001. Statistical analyses comparing mean values were performed using one of the following tests: the Friedman test with a Dunn’s multiple comparison test, the Kruskal-Wallis test with a Dunn’s multiple comparison test, or a Ordinary one-way ANOVA followed by Dunnett’s multiple comparison test. Simple comparisons were performed using an unpaired t-test. Data were cleaned by removing the highest and lowest values.

## Author contributions

Conceptualization: L.M., S.P., F.M.; Methodology: L.M., S.P., N.F., M.G., E.M., F.M.; Validation: L.M., S.P., F.M.; Formal analysis: L.M., S.P., F.M.; Data analysis: L.M., S.P., F.M.; Writing: L.M., S.P., F.M.; Supervision: F.M.; Project administration: F.M.; Funding acquisition: F.M.

## Acknowledgments

This research was supported by ATIP-Avenir program, Inserm, the Région Occitanie, ANR (ANR-21-CE17-0061, TeFiCoPa; ANR-23-CE14-0036-02, INNERCOR), FRM (REP202110014140), Support for research: I-SITE 2024 - program of excellence of the University of Montpellier, CBS2 Doctoral School, the Fondation Groupama. The RT-qPCR service was provided by the Dr Eric Jacquet at the QPCR plateform from Réseau des plateformes de Génomique Paris-Saclay at the Institut de Chimie des Substances Naturelles at Gif-sur-Yvette. We thank the MRI-DBS imaging facility, member of the France-BioImaging national infrastructure supported by the French National Research Agency (ANR-10-INBS-04, «Investments for the future»). We thank the personel of the INM animal core facility, member of the Animal Facility Network in Montpellier (RAM).

